# Division stochasticity can be transmitted to protein expression through chromosome replication

**DOI:** 10.1101/2020.09.29.319442

**Authors:** César Augusto Nieto Acuña, César Augusto Vargas García, Abhyudai Singh, Juan Manuel Pedraza

## Abstract

Stochastic fluctuations (noise) are a fundamental characteristic of protein production. Some sources of this stochasticity are still under debate. In this work, we explore how these fluctuations can originate from the stochasticity on division events. To do that, we consider the classical gene expression model with chromosome replication following the known Helmstetter & Cooper model. This model predicts intervals of the cell cycle where bacteria can have more than one copy of a particular gene. Considering the transcription rate as proportional to the number of chromosomes and division based on a continuous rate model, we explore how stochasticity in division or equivalently in cell size, could be transmitted to gene expression. Our simulations suggest that division can be an important source of such fluctuations only if chromosomes are replicating, otherwise, this noise is not well transmitted. This effect happens even if replication is deterministic. This work can be helpful for understanding cell cycle dynamics and their interplay with phenotypic variability.

## 1. INTRODUCTION

Stochasticity is a fundamental property of biochemical reactions (Elowitz et al. (2002); Blake et al. (2003)). In bacteria, this property can cause fluctuations in protein concentration with consequences for phenotype variability (Kaern et al. (2005)), physiology (Bressloff and Newby (2013)), sensing (Yi et al. (2000)) and self-organization (Misteli (2001)). Gene expression is particularly noisy (Elowitz et al. (2002)). It is known that part of the observed fluctuations in protein synthesis are intrinsic to the transcription and translation processes (Elowitz et al. (2002)). These intrinsic fluctuations arise mainly due to mRNA number variation since, unlike the proteins which have typically numbers in the thousands, mRNAs are present in low numbers inside the bacteria (typically of the order of ten) (Paulsson (2005)). Once protein is produced at rate proportional to the number of mRNA inside the cells, a variation in this RNA number—from ten to nine, for instance— is relatively high (almost 10% in some cases) resulting on a similar variation in protein synthesis. This effect is known as *noise transmission* because although intrinsic protein production is not so noisy (fluctuations in protein are about one over one thousand) these fluctuations come from the mRNA number fluctuations.

Other sources of noise are *extrinsic* and their nature is still under debate. This kind of noise is transmitted to all of the genetic pathways in the cell. This noise could be due to RNA polymerase number fluctuations, nucleotide availability and aminoacid concentration fluctuations, ribosome number fluctuations, among others. Some authors have pointed out division stochasticity as one of the possible causes of this noise (Amir (2014); Modi et al. (2017)) but we do not have a precise model which explicitly shows how stochasticity in division can transmit noise to gene expression.

In this work, we study how stochasticity in division can be transmitted to protein fluctuations. To do that we consider classical models of transcription and translation together with a Helmstetter & Cooper model for chromosome replication (Wallden et al. (2016); Levin and Taheri-Araghi (2019)). Division was modeled using a continuous rate model previously studied by us (Nieto-Acuna et al. (2019)) consisting on division triggered by the occurrence of a fixed number of successive processes each with occurrence rate proportional to the size.

First, to study the noise transmission (noise is measured by the squared coefficient of variation *CV* ^2^, which is the variance over the mean squared), we study how much noise in protein concentration can be due to chromosome replication and division. To do that we simulated the protein synthesis including noise selectively in each step of gene expression (division, replication, transcription and translation) for a simple genetic pathway with typical parameters. We show that division can be an important source of noise in gene expression.

To explore specifically how noise in division can be transmitted, we use the fact that the fluctuations on division are approximately inversely proportional to number of the needed processes (steps) to divide. Then, changing this number and, therefore, changing the division noise, we explore two scenarios: one with fixed number of chromosomes and another with dynamic chromosome replication. We observe how the noise in protein concentration is affected by the tuning of noise in size. We found that the noise in division is transmitted only in the case of dynamic replicating chromosome.

## 2. METHODS

Gene expression is taken as a discrete stochastic process modeled by a continuous-time Markov chain. The processes involved in gene expression that are considered in this model are shown in Fig. 1

**Fig. 1.**
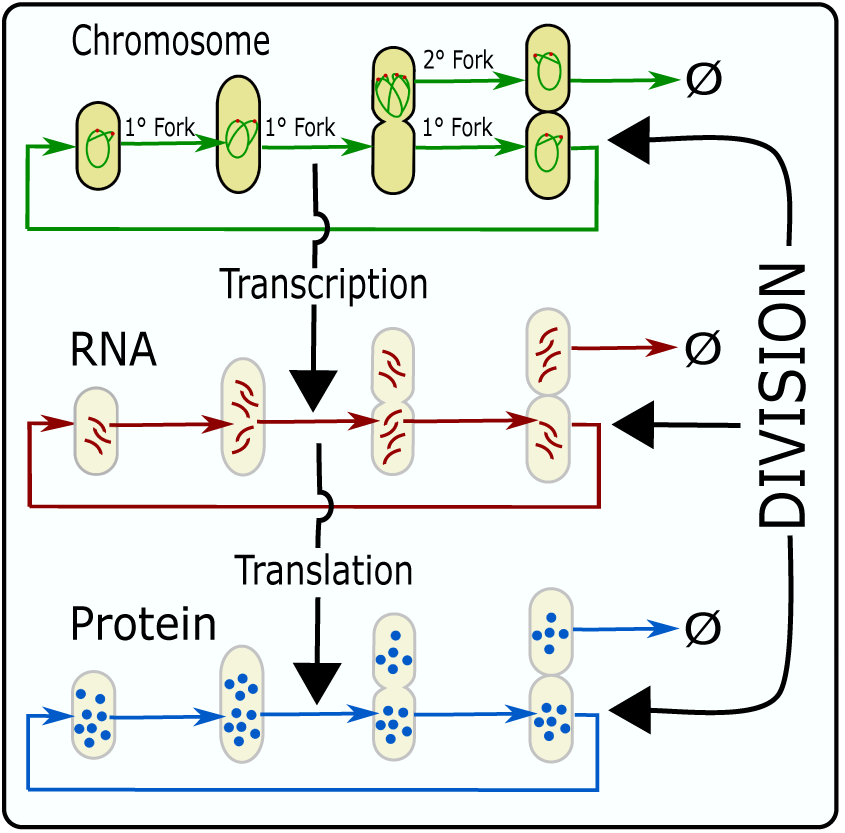
Diagram of the processes involved on gene expression.

### 2.1 Division

As we explained in previous publications (Nieto-Acuna et al. (2019); Vargas-Garc í a and Singh (2018)), division can be modeled by a continuous rate model. This consist on the occurrence of *M* steps for triggering the division. Each step can be related to the accumulation of a molecule (such as FtsZ to form the septum ring) and has a stochastic occurrence with continuous rate proportional to the size (*s*). If one supposes the growth to be exponential on time with growth rate *µ*, the associated deterministic equation describing the number of steps needed for triggering the division is

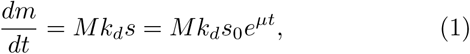

with *k*_*d*_ a constant defining the division rate. Once the *M* steps are done, division occurs and the number of steps is reset form *M* to zero and the size is halved.

By this model, the added size Δ, defined as the difference (Δ = *s*_*d*_ − *s*_*b*_) between size at division *s*_*b*_ and size (*s*_*b*_) at the beginning of the cell cycle (birth), follows an Erlang distribution:

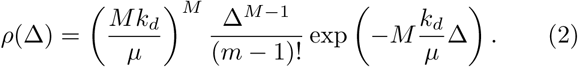

Thus, if the size at *t* = 0 is *s*_0_ and *n* divisions happened until *t*, the size at that time is given by:

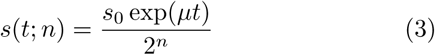

where we have considered perfectly symmetric division.

Thus, if *P*_*n,m*_(*t*) is the distribution defining the state with *n* divisions and *m* division steps up to time *t*, the master equation describing its dynamics is:

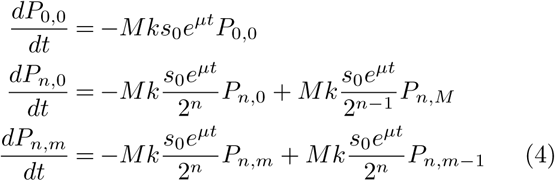

As can be observed, although there must be two sibling cells after division, in our model we track only one of these cells and the other one is discarded as is shown in Fig. 1

### 2.2 Chromosome replication

To describe chromosome replication, we use the Helmstetter & Cooper model (H & C) (Cooper and Helmstetter (1968)). As shown in Fig. 2, a replication fork starts, on the chromosome origin site, at a fixed size per origin (*s*_0_). It has been observed that this size is approximately constant over most of the growth conditions.

**Fig. 2.**
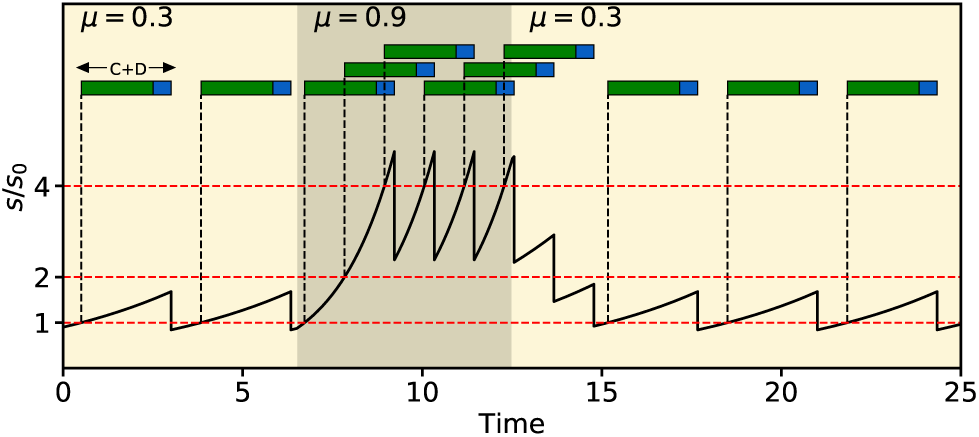
Simulation of the Helmstetter & Cooper model. In slow growth (*µ* = 0.3) only one chromosome copy is in the cell. In fast growth (*µ* = 0.9) multiple chromosome copies can exist in the cell.

During fast growth, one replication fork starts when cells reach a size *s*_0_, a second fork starts at size 2*s*_0_ and a third starts at 4*s*_0_ and so on. A relatively fixed time *C* + *D* is needed to finish the replication. Once a time *C* + *D* has passed, bacteria divide and the number of origins is halved. By this process, if bacteria grow fast multiple copies of the gene promoter can coexist inside the cell.

We model the replication as a multi-step process. The accumulation of steps 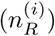 needed for replication of the *i*th fork is modeled by:

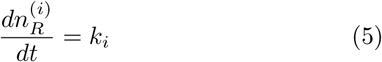

where the replication rate (*k*_*i*_) for the *i* fork is given by:

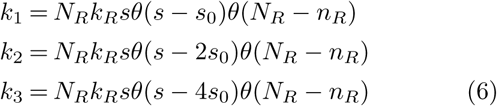

with *N*_*R*_ the number of steps needed to finish the replication, *n*_*R*_ the number of these steps that the system has already done, *k*_*R*_ a constant defining the speed of replication rate, *s* the size defined by (3), and *s*_0_ the size per origin.

The addition of the function *θ*(*N*_*R*_ − *n*_*R*_) on (6) means that replication finishes, in that fork, once the current number of replication steps *n*_*r*_ has reached the objective number of steps *N*_*r*_. The dependence of the replication rate (*k*_*i*_) on the size is considered to result in an *adder* strategy for replication. This is, adding, on average, a fixed size independently of the size at birth.

During a division event, the second fork becomes the first fork, the third fork becomes second fork and so on. The number of copies of a sample gene is chosen, arbitrarily, to duplicate every time each fork reaches half of steps needed to finish each replication.

For instance, if 20 steps are needed to finish the chromosome, the first fork has done 17 steps, the second fork has done 11 steps (one greater than the half of needed steps to finish the replication) and the third fork has done 3 steps, the number of copies of the promoter *n*_*c*_ is: one from the original chromosome, another one because the first fork has duplicated the promoter, another two due to the second fork replication and zero due to the third fork. This is, *n*_*c*_ = 4.

Although H & C considers that division happens after a period (*C* + *D*) after the replication initiation, there is evidence that division occurs in a way independent of the replication. Thus, in this work, unlike H & C, we consider the division and replication to be independent of each other.

### 2.3 Transcription

The classical model describes the concentration of RNA as a first order reaction. If the concentration of the transcription factor is fixed and its binding-unbinding rates are fast enough, the translation rate can be considered constant *k*_*r*_; if degradation is considered to happen at a rate *γ*_*r*_ and dilution by growth happens at a rate *µr*, the RNA concentration (*r*) can be modeled by the equation:

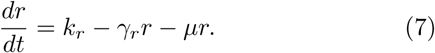

If the process is considered stochastic, usually (7) is transformed into an *Ornstein-Ulhembelck* process with *r* described as a continuous variable. However, because we are dealing mostly with discrete variables, we follow the discrete approach using master equations.

To do that, we define the number of mRNAs *n*_*r*_ inside the cell such that the concentration is given by 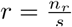 with *s* the cell size. Equation (7) can be obtained if *n*_*r*_ dynamics is given by:

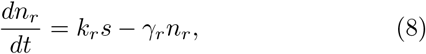

and if exponential growth 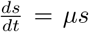 is considered and we take advantage of the properties of the derivatives:

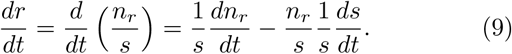

Then, a Markov chain with the number of cells inside the cell (a discrete variable) can be defined as a birth-death process with birth rate *k*_*r*_*s* as in (8) and death rate *γ*_*r*_*n*_*r*_. To obtain the mRNA concentration (*r*), one must divide this number of mRNAs (*n*_*r*_) by the current size (*s*) which is given by (3).

During division, which happens once the number of division steps *M* is reached, mRNA is divided following a binomial distribution with parameter *p* = 0.5.

In our general model, where the number of chromosomes may vary along the cell cycle, the transcription rate can be taken proportional to the number of copies of the gene promoter inside the cell. That is *k*_*r*_ → *k*_*r*_*n*_*c*_

### 2.4 Translation

Translation is, mathematically, a process similar to transcription. The dynamics of protein concentration (*p*) are given by the classical expression:

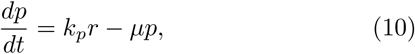

with *k*_*p*_ the translation rate per unit of *r* and degradation is usually not considered because in bacteria most proteins are not actively degraded. The term *µp* comes from dilution in a similar way to (9). In a similar way to (8), the number of proteins (*n*_*p*_) inside the cell follows:

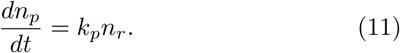

This implies that translation is a Poisson process with occurrence rate *k*_*p*_ *n*_*r*_. During division, the proteins can be approximated as splitting evenly because the number of proteins inside the cell is high (*≈*10000), so the binomial noise from this process is small.

### 2.5 Simulation of stochastic processes

Montecarlo methods are based on the frequentist approach of probability which defines an event’s probability as the limit of its relative frequency in many trials. Thus, if *n*_*t*_ is the total number of trials and *n*_*x*_ is the number of trials where the event *x* occurred, the probability *P* (*x*) of the event occurring will be approximated by the relative frequency as follows:

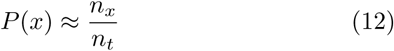

The Montecarlo approach can be summarized:

**Montecarlo approach**

1: Initialize the samples in a given state *x*_*i*_.

2: Compute the transition rates 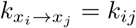

3: Define a time interval Δ*t «* (*k*_*ij*_)^*−*1^

4: Calculate the probability of occurrence of all the stochastic events during this Δ*t* by using the occurrence rates *k*_*ij*_ and integrating 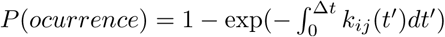

5: Generate a random number (*R*) uniformly distributed in (0,1). If this number is less than *P* (*ocurrence*), the event happens and the sample jumps from *x*_*i*_ *→ x*_*j*_.

6: Deterministic variables evolve. For instance, the time *t* = *t* + Δ*t*.

7: Repeat steps 2,3,4,5 and 6 for all the cells until the maximum time of simulation (*t* = *t*_*max*_).

8: Probability distribution is approximated by the frequentist formula (12).

When the distribution probability is approximated by equation (12), all the moments of this distribution *P* (*x*) can be estimated, particularly, the expected value 𝔼[*x*], the variance var(*x*) and the coefficient of variation *CV* ^2^(*x*):

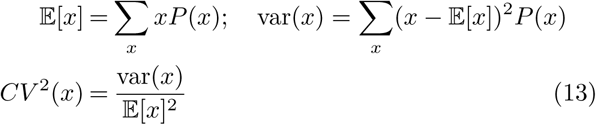

We choose typical biochemical parameters. The growth rate *µ* = 1 defines our time unit, the division rate is chosen as *k*_*d*_ = *µ*, the mean added size is 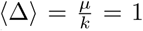 defining our unit of size and the number of division steps is chosen as *M* = 10. The mRNA degradation rate is *γ*_*r*_ = 5*µ*, and *k*_*r*_ was set such that mean concentration of mRNA was ten per cell. *k*_*p*_ was set such that the mean concentration of proteins was 1000 per cell.

Fast growth was considered. In this regime the formation of up to three forks simultaneously during replication has been observed. This can be obtained if growth rate is faster than replication rate and if the size per origin is relatively small with respect to the mean size of bacteria. Under these conditions, we set *k*_*R*_ = 0.7*µ* and 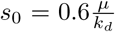. With these parameters, the average copies of the gene promoter is ⟨*n*_*c*_⟩ = 1.97. Finally, considering that replication can be a stochastic process with many steps (each step associated to a process of DNA replication) we set, arbitrarily, *N*_*R*_ = 20.

## 3. RESULTS

### 3.1 Noise transmission by the processes in gene expression

The behavior of all the variables in this work can be described by either deterministic ODEs or stochastic master equations. For instance, division has a deterministic solution using (1) or a stochastic one if (4) is used.

To determine how the total noise in gene expression can be produced by the sources considered, we sequentially set each process to be stochastic and simulate the mean noise in protein numbers over ten cycles. This is shown in Fig. 3

**Fig. 3.**
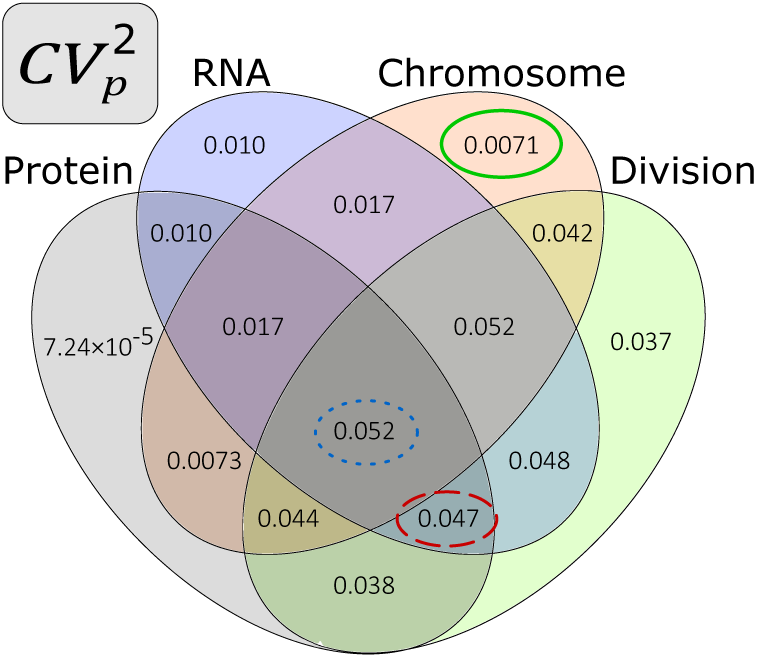
Noise in protein concentration 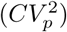 setting one or many cellular processes as stochastic. The simulation was done over 4000 cells such that the 95% confidence interval affects the significant figures after the last figure shown.

For instance, if only chromosome duplication is stochastic and the other three processes are deterministic, the resulting noise in protein concentration is 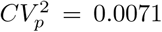, if mRNA production, division and protein synthesis are stochastic and chromosome duplication is deterministic, the noise is 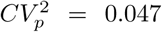. When all the sources are considered stochastic, the resulting noise is 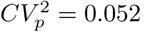.

The main result of this diagram is that when division is set as stochastic, the noise in protein concentration is relatively higher than when division is deterministic. This shows how important the division process could be as a source of noise in gene expression

### 3.2 The role of chromosome replication in transmitting division noise

To see how noise in division is transmitted, we explore three possible scenarios: stochastic replication following H & C, deterministic replication following this same model and fixed number of chromosomes per cell. This last mechanism can be interpreted as an exact replication when the cell divides such that the same number of chromosomes per cell is guaranteed. All of the parameters for the simulation were fixed to maintain the same average protein, RNA, size and chromosomes per cell.

Noise in division was modified changing the number of steps (*M*) needed to trigger division using the result from equation (2) that the noise in added size is given by:

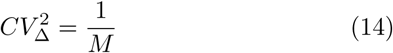

Noise in the size does not have such as simple closed expression but is known to be proportional to 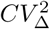.

We ran a simulation on 1000 cells for the three models changing the division steps *M* estimating the noise in both size 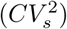 and protein concentration 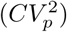 using formulas (13). The results of these simulation ares shown in Fig. 4.

**Fig. 4.**
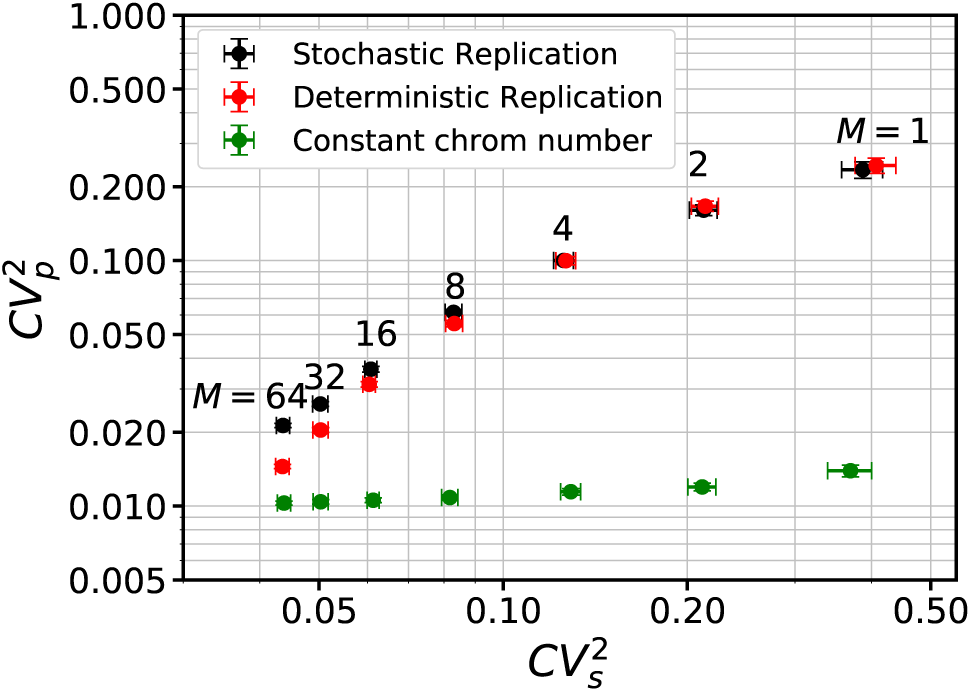
Noise of protein concentration 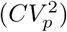 as function of the noise in size 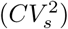 obtained changing the number of steps for division *M* from 1 to 64 in powers of 2. Error bas corresponds to the 95% confidence interval.

In Fig. 4, it can be seen how the noise in protein concentration, using the parameters listed bellow, is not very dependent on the stochasticity in chromosome replication because this 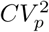 does not change appreciably when stochastic model is changed from stochastic process (black dots in Fig. 4) to deterministic one (Red dots in Fig. 4), this happens, unless the noise in division is enough suppressed (*M >* 10).

Protein noise, is highly suppressed when constant chromosome number is considered (green dots in Fig. 4) even if division is relatively noisy. In fact, the level of noise, with constant chromosome, is similar to that observed when division and chromosome are deterministic in Fig. 3 (≈0.01) as classical model expects. This effect shows how division stochasticity could be an important source of fluctuation in protein concentration mainly due to its transmission through replication process even if this is deterministic.

## 4. DISCUSSION

In this article, we have performed stochastic simulations of a model including four of the main processes involved on gene expression: Transcription, translation, chromosome duplication and cell division. We explore how much noise can is transmitted to protein concentration from each of these processes. We see that considering stochastic divisions, with a noise level similar to those found in nature, can increment the noise in protein concentration almost five times when fast chromosome replication is considered. This prevision is dramatically different to the classical model of gene expression which does not take into account the division once chromosome dynamics and division have not been described as continuous Markov chain as we consider in this work.

We also explored how this division noise is transmitted to protein concentration. To do that, we compare the noise in protein concentration changing the division noise for both a number of promoter in a replicating chromosome and a fixed number of this promoters. We observed that division noise is not transmitted to protein concentration if chromosome number does nos change along the time. This chromosome dynamics is not considered in classical models but has proven to happen experimentally (Wallden et al. (2016)).

This framework is versatile. We do not have any particular supposition of the specific nature of the chemical properties of molecules involved in all the processes. This means that this model could be used in genetically different organisms sharing similar gene expression dynamics: *E. coli* and *B. subtilis* for instance (Taheri-Araghi et al. (2015)). Adding other details to the model such as different division strategies and regulatory networks architecture can be possible for future studies bringing more features to the design of new processes involving protein production not only in bacteria but other rod-shaped organisms like yeast and archea.

Experimental Control of the the processes involved in gene expression is known to be possible (Si et al. (2018)). Translation and transcription rates can be modified by changing the chemical affinities of gene promoters ans ribosome binding sites to DNA polimerase and ribosome respectively. Division noise depends on the growth conditions and can be changed manipulating Ftsz concentration. Chromosome replication is also known to be dependent to the concentration of the replication initiator DnaA thus, an experimental study of how division noise affects the noise in gene expression through gene expression is doable.

Although chromosome replication and division noise are dependent on the growth conditions (Taheri-Araghi et al. (2015)), the effects are not well studied. For instance, in a minimal growth medium is known that division is more noisy but the number of chromosomes as seen in Fig 2, is less dynamic. This means that the noise transmitted can be very similar in both conditions: slow and fast growth. Only careful measurements can confirm our previsions.

